# Automated identification of maximal differential cell populations in flow cytometry data

**DOI:** 10.1101/837765

**Authors:** Alice Yue, Cedric Chauve, Maxwell Libbrecht, Ryan R. Brinkman

## Abstract

We introduce a new cell population score called SpecEnr (specific enrichment) and describe a method that discovers robust and accurate candidate biomarkers from flow cytometry data. Our approach identifies a new class of candidate biomarkers we define as driver cell populations, whose abundance is associated with a sample class (e.g. disease), but not as a result of a change in a related population. We show that the driver cell populations we find are also easily interpretable using a lattice-based visualization tool. Our method is implemented in the R package flowGraph, freely available on GitHub (github.com/aya49/flowGraph) and on BioConductor.

## Introduction

A major goal in flow cytometry (FCM) analysis is the identification of candidate biomarkers. The most common candidates are differential cell populations (DCPs). These are cell populations whose proportional abundances (i.e., the relative quantity of cells in a cell population) differ significantly between samples of separate classes (e.g. disease vs healthy). Commonly used metrics for proportional abundance are cells per *μ*L of blood and proportion (i.e, the ratio between the count of cells in a population and some parent population).

Here, we propose the concept of *maximal differential cell populations* (MDCPs). MD-CPs are DCPs whose change in proportional abundance is only significantly associated with its sample class, as opposed to being the result of a proportional abundance change in a related DCP. For example, if there is a significant decrease in the proportion of helper T-cells in samples from sick individuals compared to those from healthy individuals, then helper T-cells is a DCP. However, if the proportional abundance of all types of T-cells decrease at a similar rate, then we can hypothesize that the disease decreases the proportional abundance of all T-cells. It follows that T-cells and all of its child populations, including helper T-cells, are DCPs but only T-cells is a MDCP. MDCPs are preferable candidate biomarkers because their proportional abundance change is only driven by their association with a sample class. We refer to such cell populations as driver cell populations. To our knowledge, while there are many methods that find biomarker candidates by identifying DCPs, there are no methods that do so by isolating only the MDCPs among those DCPs.

Most methods identify DCPs either as a byproduct of another procedure [5, 7, 8, 18, 23] (e.g., CytoDX [11] main goal is to classify FCM samples, but it also tries to find DCPs as a postprocessing step) or compare prespecified cell populations by evaluating whether there is a large difference in their proportional abundance across samples using some statistical significance test [12, 14, 19]. A summary of related methods can be found in [1, 9]. Though there are methods that attempt to find MDCPs by expanding their DCP candidates to cell populations that are dependent on each other, the statistical tests they use assume independence between cell populations. For example, Cydar [13] uses the spatial false discovery rate, and diffcyt [21] uses statistical tests traditionally used for differential analysis in bulk RNAseq data sets where the transcripts/genes are assumed to be independent of each other. Such statistical tests would only be able find DCPs despite the large MDCP candidate pool because they do not account for the relationships between cell populations.

To address these shortcomings, we identify MDCPs by comparing the SpecEnr of cell populations across samples, where SpecEnr is a novel cell population score, a numerical metric, that is derived from the proportional abundance metric, proportion, and accounts for the relationship between cell populations.

In this paper, we

1. Define and formulate the problem of finding driver cell populations by identifying MDCPs,
2. Introduce a cell population score SpecEnr (specific enrichment) that accounts for dependencies between parent and child cell populations, and
3. Describe a method that harnesses SpecEnr properties to find robust, accurate, and easily interpretable driver cell populations.

We hypothesize that identifying MDCPs will aid the understanding of disease etiology.

## Methods and Materials

### Preprocessing

To calculate SpecEnr, we take as input a vector of cell population proportions for each FCM sample generated using any suitable manual or automated approach. For our experiments, given a FCM sample containing a cell × measurement matrix and threshold gates obtained via gating, we use flowType [14] to identify all possible cell populations and enumerate their cell count. Next, we normalize cell counts relative to the total count of cells in each sample. We do so by converting counts into proportions by taking the cell count of each cell population over the total number of cells in the sample.

Users can also choose to identify cell populations via methods other than flowType. The requirement for calculating the SpecEnr of a cell population is that its and all of its parents and grandparents proportions should be available. For example, if the user chooses to identify cell populations via clustering, we can treat each cluster as a unique gate; this way, a cell population’s parent cell populations would include all possible pairwise combinations of it and all other cell populations. Its grandparent cell populations would be all possible mergers of it and any two other cell populations.

### Cell hierarchy

To visualize the relationship between cell populations, we use the cell population hierarchy of a sample. A *cell population hierarchy* or cell hierarchy for short, is a directed acyclic graph where nodes represent cell populations and edges represent the relationship between cell populations (Figure 1). We define a cell population as a set of cells with similar fluorescent intensity (FI) values for a set of 0 ≤ *l* ≤ *L* measurements (e.g. markers, SSC, FSC). For simplicity, we define a measurement condition as a combination of a measurement and a positive^+^ or negative^−^ expression indicator. For example, *A*^+^*B*^−^ contains two measurement conditions (*A*^+^ and *B*^−^) and represents a cell population whose cells have FI greater and less than the given thresholds for measurements *A* and *B* respectively. We define the *l*’th layer of the cell hierarchy as the set of all nodes whose label contains exactly *l* unique measurement conditions. It then follows that a cell hierarchy has *L* + 1 possible layers. The 0’th layer contains the root cell population comprising all cells.

**FIGURE 1:**
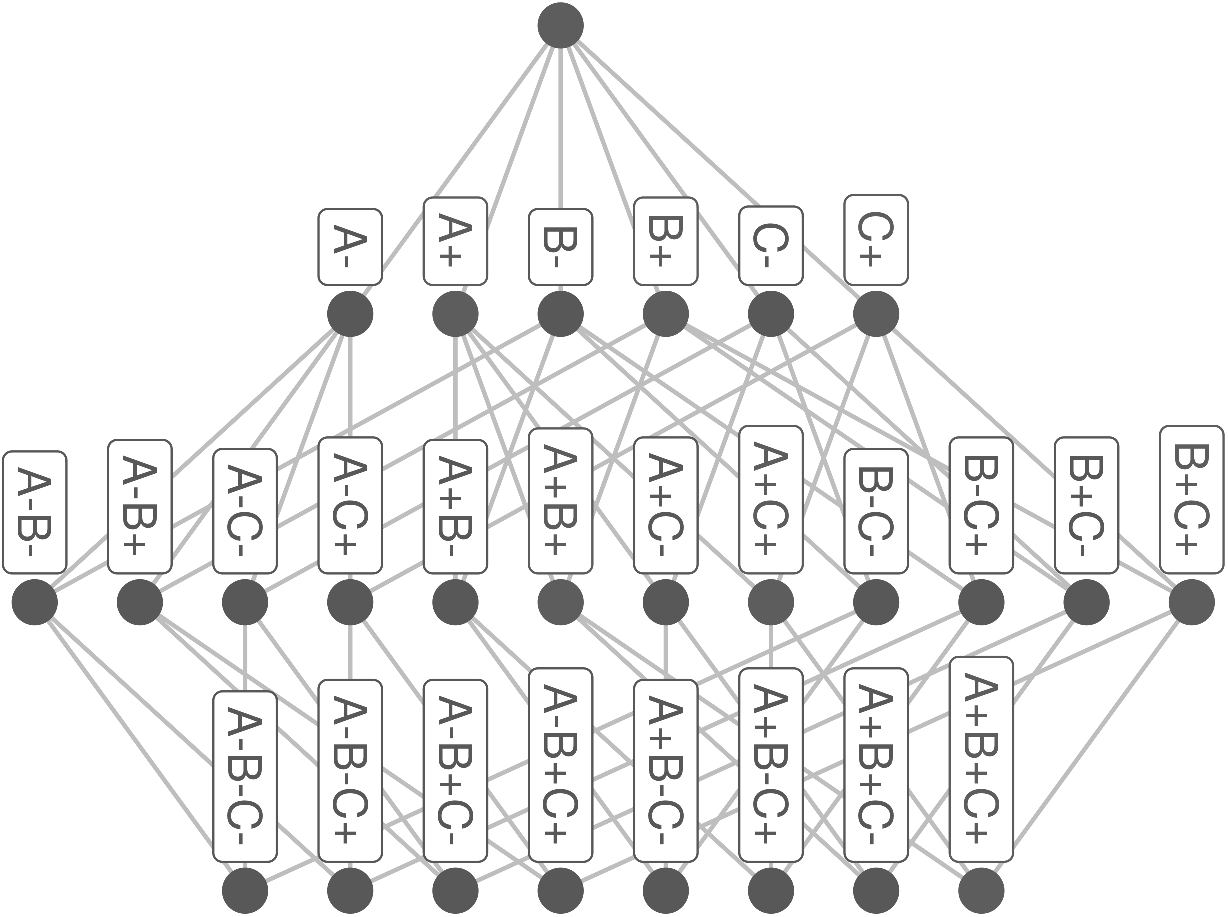
An example of a cell population hierarchy representation of a FCM sample and its cell populations defined by measurements *A*, *B*, and *C*.

In the cell hierarchy, each edge points from a ‘parent’ cell population to its ‘child’ sub-population defined by the addition of one measurement condition. For example, if there are three measurements {*A, B, C*}, then there are edges from the node representing the cell population labelled *A*^+^ to the nodes labelled *A*^+^*B*^+^, *A*^+^*B*^−^, *A*^+^*C*^+^, and *A*^+^*C*^−^.

We denote the actual proportion *P* of any node *v*^1:*l*^ in layer *l* by *P* (*v*^1:*l*^) such that 1:*l* (1, 2, …, *l*) are the indices of the measurement conditions its label contains.

We show in Supplementary Material Section 2 that we can derive our methods’ scores for all cell populations just from those cell populations whose labels only contain positive conditions. Following this reasoning, and to simplify our notation, in the following sections, we assume that the measurement conditions used to label our cell populations are all positive. This implies that the measurements used must be unique, as they always should be. For example, cell population *A*^+^*B*^+^*C*^+^ has three positive measurement conditions and can therefore be denoted as *v*^1:3^; subsequently, we can denote its parents *A*^+^*C*^+^ and *A*^+^*B*^+^ as *v*^{1:3}\22^ and *v*^{1:3}\3^ by excluding the second and third measurement conditions.

### Cell population score: SpecEnr

The assumptions we will introduce for SpecEnr are based on well-established concepts in probability theory [3]. Measurement conditions are random events, and the proportion of each cell population is the probability of jointly occurring random events.

To obtain SpecEnr, we compare the actual proportion of a cell population with its expected proportion: the proportion we expect a cell population to have given the proportion of its ancestors. By doing so, we can evaluate its proportion changes independent of the effects incurred by its ancestors.

#### Expected proportion

The SpenEnr null hypothesis imagines that each cell population has at least two measurement conditions that are independent given the others. Specifically, under the null hypothesis, for a cell population *v*^1:*l*^ with proportion *P* (*v*^1:*l*^), the following holds. Without loss of generality, let us assume that *P* (*v*^1^) (e.g. *A*^+^) and *P* (*v*^2^) (e.g. *B*^+^) are independent given *P* (*v*^3:*l*^) (e.g. *C*^−^).

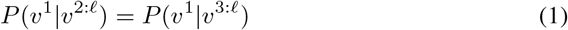

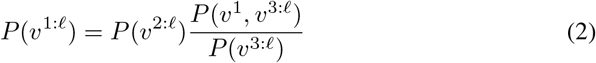

where *P* (*v*^1^|*v*^3:*l*^) indicates the conditional proportion of *v*^1^ given *v*^3:*l*^.

Generalizing this assumption to any *p, q* pair, *p* ∈ 1:*l* and *q* ∈ 1:*l* \ *p*, we get

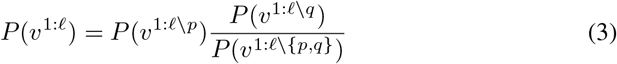

While this assumption may be applied to most cell populations, there are edge cases. Our assumption requires *P* (*v*^1:*l\{p,q}*^) to exist. Therefore, expected proportion is only calculated for cell populations in layers *l* ≥ 2. For the root node, we initialize its expected proportion to 1. For the nodes in layer one, we initialize their expected proportions to. By initializing their expected proportion to 0.5, we maintain the sum-to-1 rule in probability where, for example, *P* (*A*^+^) + *P* (*A*^−^) = 1.

To identify differential cell populations, we compare their expected and actual proportion. In Equation 3, we assumed all measurement condition pairs, with indices {*q, p*}, *P* (*v^p^*) and *P* (*v^q^*) to be independent give *P* (*v*^1:*l\{p,q}*^). Now let us assume that this does not hold for *A*^+^*B*^+^*C*^+^’s parent cell population *A*^+^*C*^+^. While *A*^+^ and *C*^+^ are dependent on each other, *B*^+^ is independent of both *A*^+^ and *C*^+^. In this case, the assumption we made in Equation 2 only holds for cell population *A*^+^*B*^+^*C*^+^ when *q* ∈ {1, 2} and *p* = 3. We do not want to flag *A*^+^*B*^+^*C*^+^ as maximally differential as its proportion change is completely dependent on cell populations *A*^+^*C*^+^ and *B*^+^. Therefore, we relax our assumption in Equation 3 to: there must be some index pair {*p, q*} such that *P* (*v^p^*) is independent of *P* (*v^q^*) given *P* (*v*^1:*l\{p,q}*^). Then *P* (*v*^1:*l*^) can be calculated as follows.

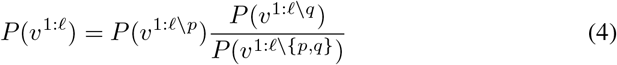

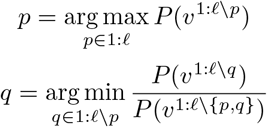

Otherwise, if there is no *p, q* pair such that *P* (*v^p^*) is independent of *P* (*v^q^*), then Equation 4 does not hold and *P* (*v*)’s abundance change cannot be attributed to any of its ancestors’ abundance change.

Additional details on proof of correctness for our assumption is in Supplementary Material Section 1.

#### SpecEnr

In this section, we explain how we calculate our proposed SpecEnr score. Given the expected proportion of cell population *v* calculated using Equation 4, SpecEnr is the natural log of *v*’s actual proportion over its expected proportion calculated using Equation 4 which we denote here as *E*(*v*).

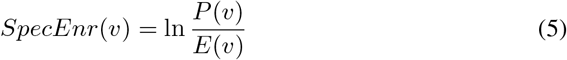

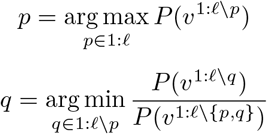

SpecEnr accounts for the dependency of a cell population on its ancestors. For example, if a cell population has a SpecEnr value of 0, then its proportional abundance is completely dependent on that of its ancestors. Otherwise, it contains measurement conditions that are all dependent on each other, where *P* (*v^p^*) is dependent on *P* (*v^q^*) for all {*p, q*} ∈ 1:*l* (i.e. Equation 3 does not hold for any *p, q*).

The asymptotic runtime and actual runtime to calculate SpecEnr are provided in the Supplementary Material Sections 2 and 7.

#### Maximal differential cell population (MDCP)

A maximal differential cell population (MDCP) is a cell population that has significantly different abundance across sample groups. In this respect, a MDCP is similar to a differential cell population (DCP). However, in addition to this, MDCP have an additional property where its abundance difference cannot solely be attributed to the abundance difference in its ancestor cell populations. Therefore, a MDCP’s abundance change is unique and can help users confirm or reject hypotheses in biological experiments.

SpecEnr is an example of log-probability ratios, which are commonly used in Bayesian hypothesis testing [6]. Following a similar framework, a cell population is not a MDCP if its SpecEnr values across classes are not significantly different. Conversely, in order for a cell population to be a MDCP *v*^1:*l*^, it must satisfy two conditions.

1. A MDCP SpecEnr must be significantly different between samples according to a filtered adjusted T-test we describe in the next section.
2. A MDCP must also be maximal, in that it must not have any direct descendants who meet the first condition above.

The second condition is required because our first is also satisfied by direct ancestors of a MDCP as its ancestor cell populations are defined by a subset of measurement conditions defining the MDCP.

Relating our definition back to the difference between MDCP and DCP : if there exists one MDCP in our data set (e.g. A^+^B^−^), then the DCPs would be the MDCPs’ ancestors (e.g. A^+^ and B^−^) and descendants (e.g. A^+^B^−^C^+^ and A^+^B^−^C^−^), and all cell populations that share at least one measurement condition with the MDCP (e.g. B^−^C^+^ and A^+^C^−^). This further demonstrates the difficulty of identifying MDCPs among DCPs; as all DCPs would be a candidate MDCP.

### Testing whether the cell population SpecEnr values across sample classes are significantly different

To test if a cell population satisfies our first condition for MDCPs (i.e. its SpecEnr is significantly different across samples), we apply the T-test on SpecEnr values for each cell population across two sets of samples (e.g. a control group and an experiment group). We show that the raw SpecEnr T-test p-values are statistically sound in Supplementary Material Section 5.

Given that we are testing multiple hypotheses, we adjust the p-values *ρ_v_* for each cell population *v* using layer-stratified Bonferroni correction [2] to obtain our final adjusted p-values *ρ*′_*v*_. We do so by multiplying our p-values with the number of cell populations in the layer on which cell population *v* resides *m_l_* and the total number of layers *L* + 1 (including the layer 0; see Supplementary Material Section 3 for additional details).

We use a q-value (i.e., the adjusted p-value) threshold < .05 to determine if a cell population q-value is significant and potentially maximally differential.

#### Avoiding falsely significant q-values with filters

In some cases, the p-value obtained by evaluating SpecEnr may be falsely significant when dealing with small or noisy data sets. As a cell populations proportion gets close to 0, the actual versus expected proportion ratio used to calculate SpecEnr becomes inflated. As well, if we are conducting significance tests on cell populations with SpecEnr values of 0 (i.e. actual and expected proportions are the same) model-based significance tests (e.g. T-test) are highly influenced by outliers and rank-based significant tests (e.g. Wilcoxon) are influenced by random ordering of 0’s. To ensure our SpecEnr p-values are valid, we mark cell populations as insignificant if any of the following apply.

1. They do not have a mean count of a user-specified threshold of events (we use > 50 for our data sets) to prevent inflated ratios,
2. They do not have significantly different actual versus expected proportions for at least one of the sample classes, and
3. They have actual and expected proportions that are significantly different across both sample classes.

In our experiments, we use a standard significance threshold of < .05 for all T-test p-values on filter related significance tests. We show an example of these filters in the Supplementary Material Section 4.

For brevity, we call the p- and q-values obtained using SpecEnr and proportion, SpecEnr p- and q-values, and proportion p- and q-values respectively.

### Experimental data

To confirm that flowGraph is able to identify known MDCPs we prepared synthetic negative and positive control data sets and used two previously published biological data sets.

#### Synthetic data

- **neg1** (Negative control): For each cell, we assigned it to be positive^+^ for each measurement with a 50% probability.

- Samples: 10 control vs 10 experiment (300,000 cells/sample).
- Measurements: *A*, *B*, *C*, and *D*.
- **pos1** (Positive control 1): Same as neg1, except in the experiment samples, cell population *A*^+^ is increased by 50%. More specifically, in each *R* × *L* matrix, we duplicated a random sample of half the cells in *A*^+^.
- **pos2** (Positive control 2): Same as pos1, except instead of *A*^+^, *A*^+^*B*^+^*C*^+^ is increased by 50%.
- **pos3** (Positive control 3): Same as pos1, except instead of *A*^+^, a random sample of half of all cells that belong to at least one of *A*^+^*B*^+^ and *D*^+^ are duplicated (i.e. increased by 50%), indirectly causing a unique increase in cell population *A*^+^*B*^+^*D*^+^. Note that cells that belong to both *A*^+^*B*^+^ and *D*^+^ are duplicated once instead of twice to ensure both cell populations increase by 50%.

#### Real data sets

- **flowcap** (FlowCAP-II AML data set): This data set is from the FlowCAP-II [1], AML challenge, panel 6. It is known that AML samples have a larger *CD34*^+^ population [1].

- Samples: 316 healthy vs 43 AML positive subjects’ blood or bone marrow tissue samples (~60,000 cells/sample).
- Measurements: *HLA-DR*, *CD117*, *CD45*, *CD34*, and *CD38*.
- **pregnancy** (Immune clock of pregnancy data set): While the previous two data sets are flow cytometry data sets, this is a CyTOF data set. Nevertheless, our method can be used on either type of data set. So far, there has been no experiments that identified ground truth driver cell populations for the pregnancy data set [4]. However, the original authors were able to train classifiers on the same patients using FCM and multi-omics data [15]. Therefore, we hypothesize that we will be able to find MDCPs in this data set that are associated with the sample classes listed below.

- Samples: 28 late-term pregnancy vs 28 6-weeks postpartum human maternal whole-blood samples (~300,000 cells/sample); Samples are taken from each of the 18 and 10 women of the training and validation cohort during late-term pregnancy and 6 weeks postpartum.
- Measurements: *CD123*, *CD14*, *CD16*, *CD3*, *CD4*, *CD45*, *CD45RA*, *CD56*, *CD66*, *CD7*, *CD8*, *Tbet*, and *TCRgd*.
- To account for possible batch effects associated with the subjects who provided the FCM samples, we used the paired T-test where samples were paired with respect to subject.

## Results

### SpecEnr p-values are robust

We hypothesized that theoretically similar data sets yield similar unadjusted p-values across all cell populations. To test this, we split up the samples in data set pos1 in half and compare the samples across these two halves or “theoretically similar data sets”. When we compared the unadjusted SpecEnr p-values across these theoretically similar data sets using the Spearman correlation, we obtained a perfect score of 1. We saw the same result with metrics recall, precision, and F measure over the first set. These results indicate that significant cell populations in the first set also show up as significant in the second set. We also show that unadjusted SpecEnr p-values are statistically sound with the following experimental results [20]. Using SpecEnr, we were able to generate a random uniform distribution of unadjusted p-values on our negative control data set neg1. It follows that 5% of the SpecEnr p-values were below our .05 threshold (See Supplementary Material Section 5 for added detail).

### SpecEnr q-values help identify accurate driver cell populations in synthetic data sets

flowGraph accurately identified that pos1 and pos2 driver cell populations were *A*^+^ and *A*^+^*B*^+^*C*^+^ (Figure 2). While both SpecEnr and proportion q-values flagged these cell populations, when we observed SpecEnr q-values, the descendants of these driver cell populations were not flagged as significant. This was also true when multiple driver cell populations were present in lower layers of the cell hierarchy. In pos3, where both *A*^+^*B*^+^ and *D*^+^ were increased to cause a unique change in *A*^+^*B*^+^*D*^+^; we saw that SpecEnr q-values were only significant for these three cell populations and their ancestors. Results from our positive control data sets were also similar when the same cell populations decreased instead of increased in proportional abundance (Supplementary Material Section 6).

**FIGURE 2:**
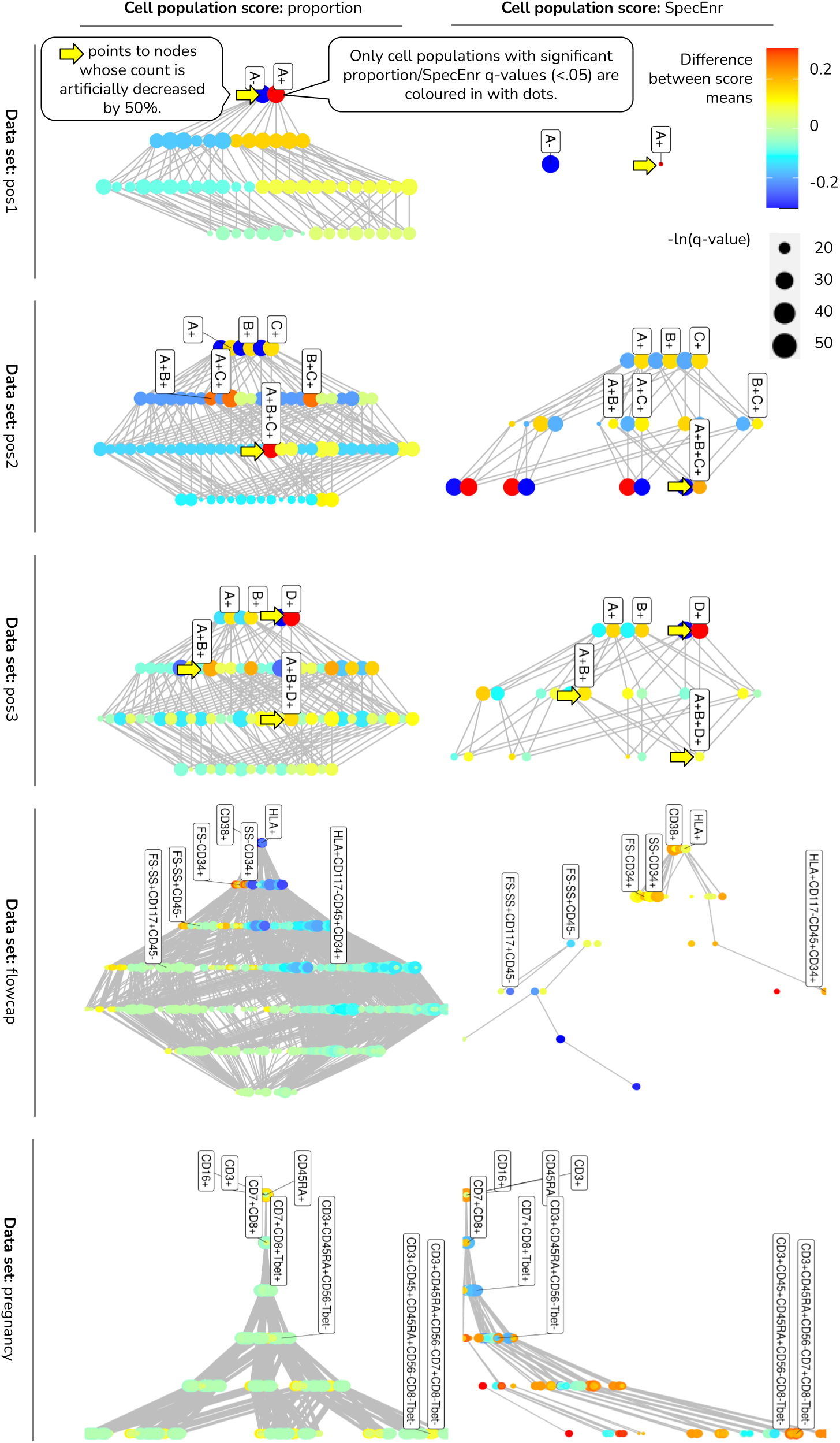
Cell hierarchy plots for synthetic data sets pos1-3 and real data sets flowcap and pregnancy show that SpecEnr q-values accurately identify MDCPs while proportion q-values flag all DCPs but do not highlight which of the DCPs are MDCPs.

### SpecEnr q-values flag known and novel driver cell populations in real data sets

For the flowcap data set, SpecEnr directs users down a branch of the cell hierarchy from physical properties *SS*^+^ and *FS*^−^ to *FS^−^SS*^+^*CD117*^+^*45*^+^ and *HLA*^+^*CD117^−^CD45*^+^*CD34*^+^. While *HLA* and *CD117* are variably expressed on cells in FCM samples from subjects with AML [16, 22], *CD34* and *CD45* are expressed on blast cells [10, 17]. This is important as the abundance of blast cells aid in diagnosis of AML [1].

In the pregnancy data set, the top most significant cell populations displayed by our statistical significance test showed an up-regulation in cell populations containing *CD3*, *CD45*, and *CD45RA* (e.g. *CD3*^+^*CD45RA*^+^*CD56^−^Tbet^−^*). SpecEnr q-values also indicate that cell populations containing measurements *CD8* and *CD16* are significantly down-regulated. Meanwhile, proportion q-values flag all DCPs in the cell hierarchy as significant.

## Discussion

In this paper, we introduced a new cell population score, SpecEnr, and a method, flow-Graph, that integrates SpecEnr to identify MDCPs. We showed that the results of flow-Graph are statistically sound, accurate, and easily interpretable.

In the FlowCAP-II challenge, the AML data set was used to evaluate how well methods are able to classify samples belonging to healthy and AML positive subjects. Among the competing methods, those that used cell population proportions for classification were DREAM–D, flowPeakssvm, Kmeanssvm, flowType, FeaLect, PBSC, BCB/SPADE, SWIFT [1]. All of these methods assume that cell count and proportion may be used to differentiate between the two classes of samples. However, cell count and proportion do not account for relations between cell populations, making it difficult to isolate the MDCPs among the DCPs (Figure 2). To account for these relationship, one can manually analyze the ratio of the count of cells in a population over all of its direct parent populations. However, given *L* measurements, there are 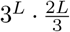 such relationships not including the relationship between a cell population and its indirect ancestors [14]. In contrast to comparing 3*^L^* cell population scores, directly comparing cell population relations becomes computationally impractical.

SpecEnr mitigates both problems as it is a cell population score that accounts for relations between cell populations. q-values obtained from computed SpecEnr scores isolated only the few ground truth driver cell populations (MDCP e.g. *SS^−^CD34*^+^). Our results not only reveal known driver cell population *CD34*^+^ but also provide visualizations signifying that their change was caused by a change in its descendants exposing novel driver cell populations.

We also observed this contrast in behaviour between SpecEnr and proportion q-values in the pregnancy data set. We hypothesized that flowGraph w ould be able to find MDCPs because [4] were able to use L_1_, L_2_, and cell signal pathway regularized regression to classify samples taken from women at different stages of pregnancy. The original authors also assumed that there exists MDCPs in the pregnancy data set [11]. However, because these methods find candidate biomarkers as a byproduct of a sample classification method, there was no way of verifying whether the candidate biomarkers they inferred are simply DCPs or are also MDCPs. FlowGraph answers this question by providing users a way to differentiate between the two while verifying our hypothesis validating the existence of MDCPs in the pregnancy data set.

### Future work

Since SpecEnr is calculated using proportions, it is prone to the same issue that occur when using proportions directly. That is, changes in proportion of cell populations must sum to 0. For example, in pos1, *A*^+^ abundance doubled, so its proportion increased from .5 to .66; but *A*^−^ proportion decreased from .5 to .33. More generally, if a cell population is differential, it will induce a change in the proportion of all cell populations that are labelled using the same set of measurements as it; because these cell populations are mutually exclusive. Another example of this are the {*A*^{+,−}^*B*^{+,−}^*C*^{+,−}^} cell populations from pos2. If the driver cell population resides in layers > 1, then it is easily identifiable as the cell population with the largest magnitude of change. In the future, we would like to improve on our method such that we only flag the driver cell populations and not the cell populations it affects in the context of proportions.

Finally, we showed that an adjusted and filtered T-test on SpecEnr will yield a significant q-value on driver cell populations and their ancestors. While this makes driver cell populations intuitive to find on a cell hierarchy plot, ideally, we should only flag the driver cell populations as significant and not their ancestors. By preventing excessive flagging of ancestor populations, we enable more expressive and detailed insights in results interpretation.

## Acknowledgements

This work was funded by Simon Fraser University and the Natural Sciences and Engineering Research Council (RGPIN-2020-04903).

