## Supplementary Material for "Automated identification of maximal differential cell populations in flow cytometry data"

Alice Yue, Cedric Cauve, Maxwell Libbrecht, Ryan R. Brinkman

#### 1 Proof of correctness for Equation 4.

In this section, we work under the assumption that there exists some pair of measurement conditions that are independent of each other.

**Theorem 1.1.** *Assuming that there exists some pair of measurement condition indices* $\{p, q\}$ *s.t.* $P(v^p)$ *and* $P(v^q)$ *are independent given* $P(v^{1:\ell\setminus\{p,q\}})$*:*

$$P(v^{1:\ell}) = P(v^{1:\ell\setminus p})\frac{P(v^{1:\ell\setminus q})}{P(v^{1:\ell\setminus\{p,q\}})}$$

*where* $P$ *is the actual proportion of cell population* $v$ *defined by* $\ell$ *measurement conditions, then we can identify such an index pair using the following.*

$$p = \arg\max_{p\in 1:\ell} P(v^{1:\ell\setminus p})$$

$$q = \arg\min_{q\in 1:\ell\setminus p} \frac{P(v^{1:\ell\setminus q})}{P(v^{1:\ell\setminus\{p,q\}})}$$

*Proof.* In our method, we derived equation 4 by assuming there is some $\{p, q\}$ index pair s.t. $P(v^p)$ is independent of $P(v^q)$ given $P(v^{1:\ell\setminus\{p,q\}})$. For purposes of this proof, we will use an alternative set of notations:

$$\sum_z p(z) = P(v^{1:\ell\setminus\{p,q\}}) + (1 - P(v^{1:\ell\setminus\{p,q\}})) = 1$$

$$\sum_x p(x) = P(v^{1:\ell\setminus p}) + (1 - P(v^{1:\ell\setminus p})) = 1$$

$$\sum_y p(y) = P(v^{1:\ell\setminus q}) + (1 - P(v^{1:\ell\setminus q})) = 1$$

where $p$ is the probability distribution function. In our application, each cell is either a part of the our cell population $v$, or it is not. This means that our probability distributions $p$ are all binary and add up to 1.

First show the relationship between conditional mutual information and Equation 4.

$$
\begin{aligned}
I(X;Y|Z) &= \sum_{x,y,z} p(x,y,z) \log \frac{p(x,y|z)}{p(x|z)p(y|z)} \\
&= \sum_{x,y,z} p(x,y,z) \log \frac{p(x,y,z)p(z)}{p(x,z)p(y,z)} \\
&= \sum_{x,y,z} p(x,y,z) \left( \log p(x,y,z) + \log \frac{p(z)}{p(x,z)p(y,z)} \right) \\
&= \sum_{x,y,z} p(x,y,z) \log p(x,y,z) + \sum_{x,y,z} p(x,y,z) \log \frac{p(z)}{p(x,z)p(y,z)}
\end{aligned}
$$

Conditional mutual information $I(X;Y|Z) = 0$ iff $X$ is independent of $Y$ given $Z$. We assumed such a pair exists. However, even if we are not able to find such $X$ and $Y$, we also know that conditional mutual information must always non-negative $I(x;Y|Z) \geq 0$. Given these two properties, the following is true.

$$
\begin{aligned}
0 &\leq \sum_{x,y,z} p(x,y,z) \log p(x,y,z) + \sum_{x,y,z} p(x,y,z) \log \frac{p(z)}{p(x,z)p(y,z)} \\
-\sum_{x,y,z} p(x,y,z) \log p(x,y,z) &\geq \sum_{x,y,z} p(x,y,z) \log \frac{p(z)}{p(x,z)p(y,z)} \\
\sum_{x,y,z} p(x,y,z) \log p(x,y,z) &\leq \sum_{x,y,z} p(x,y,z) \log \frac{p(x,z)p(y,z)}{p(z)}
\end{aligned}
$$

In order to find a $X$ and $Y$ such that they are independent given $Z$, we must minimize the difference, to 0, between two sides of this inequality. In order to do so, we must choose {p,q} according to Equation 4, concluding our proof.

□

### 2 Algorithm: calculating SpecEnr values for each cell population given proportions.

To reduce runtime, we only directly calculate this expected proportion for cell populations with only positive measurement conditions (e.g. $A^+B^+$, $C^+D^+$); the expected proportion for the rest of the cell populations can be directly inferred from these cell populations in a later step. This runtime includes operations to first calculate all the arc values in our cell hierarchy. Arc values can be interpreted as the proportion of the cell population an arc points to over the proportion of the cell population an arc originates from. Since there are $\sum_{\ell=0}^{L} \left\{ l \cdot \begin{pmatrix} L \\ \ell \end{pmatrix} \right\} = 2^{L-1}L$ edges, this step takes $O(2^L)$ operations. Next, to find the maximum parent proportion and minimum value on edges pointing to the non-maximum parent, we do $\ell$ and $\ell(\ell-1)$ operations for each cell population. Given that there are $\begin{pmatrix} L \\ l \end{pmatrix}$ cell populations with only positive measurement conditions on each layer, this part requires $\sum_{\ell=2}^{L} \ell^2 \begin{pmatrix} L \\ l \end{pmatrix}$ or $O(\ell^3)$ operations. If we were to calculate the above for all cell populations, the first part would require $O(3^L)$ as opposed to $O(2^L)$ operations and we would have to consider $2^\ell \begin{pmatrix} L \\ l \end{pmatrix}$ instead of $\begin{pmatrix} L \\ l \end{pmatrix}$ cell populations per layer for the second part.

To follow up on all other cell populations, we observe that the score values for these cell populations collectively implicitly contain information about all nodes in the full hierarchy. As such, we can deduce the expected proportion of all other cell populations with at least one negative measurement condition. For example, the first layer of a cell hierarchy can be specified with $L$ cell populations. The count of these cell populations with a negative measurement condition (e.g. $A^-$) can be deduced by the difference between the total proportion and the proportion of the corresponding cell population expressing positively for that measurement (e.g. $P(A^-) = P(root) - P(A^+)$). The $L$'th layer can be specified with one cell population i.e. a cell population with all measurements. The total number of cell populations needed to specify all cell populations is $2^L$ or the number of cell populations with only positive measurement conditions.

**Data:** $V \leftarrow$ cell populations with all positive measurement conditions whose expected proportion is already calculated.
**Data:** $V^- \leftarrow$ all cell populations with $\geq 1$ negative measurement condition(s).
**for** $\ell := 2 \rightarrow L$ **do**
  $\mathbf{v}_\ell \leftarrow \{v | v \in V, |v| = \ell\}$ **while** $v_\ell \neq \{\}$ **do**
    $\mathbf{v}^*_\ell \leftarrow \{\}$
    **for** $v_\ell := \mathbf{v}_\ell$ **do**
      **for** $p^+ :=$ a measurement in $v_\ell$ with a positive$^+$ condition **do**
        $P(v^{p^-,1:\ell \setminus p^+}) = P(v^{1:\ell \setminus p^+}) - P'(v_\ell)$
        $\mathbf{v}^*_\ell \leftarrow \{\mathbf{v}^*_\ell, v^{p^-,1:\ell \setminus p^+}\}$
        $V = \{V, v^{p^-,1:\ell \setminus p^+}\}$
        $V^- = \{V^- \setminus v^{p^-,1:\ell \setminus p^+}\}$
      **end**
    **end**
    $\mathbf{v}_\ell = \mathbf{v}^*_\ell$
  **end**
**end**

**Algorithm 1:** Calculating the expected proportion of cell populations with negative measurement conditions

From this, we use Algorithm 1 to infer the expected proportion of all other cell populations with at least one negative measurement condition. This algorithm takes one difference operation per cell population amounting to $O(3^L)$ operations total. Since we only calculate expected proportions for $O(2^L)$ cell populations and since the total number of other cell populations far out-scales $O(2^L)$, we achieve an overall runtime of $O(3^L)$.

In the case that there is more than one threshold per measurement (e.g. on top of $A^-$ and $A^-$, there also exists $A^{++}$, $A^{+++}$, and so on), the proof of correctness for expected proportions still holds as a cell population in layer $\ell$ must have $\ell$ parent nodes. The correctness of algorithm 1 also still holds except the proportions represented by index $p+$ is now not a single proportion value but a sum of all proportions with a positive measurement condition for the measurement on index $p$. Note that this is contingent on a characteristic of flowType who labels measurements with additional thresholds as positive measurement conditions. Conversely, there can be only one negative measurement condition per measurements (i.e. there can be no $A^{--}$). As we still only calculate expected proportion for cell populations containing exclusively positive measurement conditions, the runtime of expected proportion calculation increases with the number of thresholds. If each measurement is assigned $k \geq 1$ thresholds, then the number of cell population containing only positive measurement conditions is $O((k+1)^L)$ while the total number of cell populations is $O((k+2)^L)$. The overall runtime then becomes $O((k+2)^L)$.

#### 3 Layer-stratified Bonferroni correction

We define layer-stratified Bonferroni correction as follows. Let $p_{1:m}$ be p-values associated with a set of hypotheses $H_{1:m}$. For $\ell \in 1 \ldots L$, let $\sigma_\ell$ be a subset of hypotheses $\{1 \ldots m\}$ (e.g. cell populations in particular layer of a hierarchy). For $i \in \sigma_\ell$, we define the adjusted p-value $p'_i = p_i|\sigma_\ell|L$. We accept a hypothesis $H_i$ if $p'_i < \alpha$.

**Theorem 3.1.** *The layer-stratified Bonferroni correction has a family-wise error rate (FWER) of at most $\alpha$.*

*Proof.*

$$
\begin{aligned}
& P\left(\bigcup_{i=1}^{m}(p'_i \leq \alpha)\right) \\
&= P\left(\bigcup_{i=1}^{m}(p_i|\sigma_\ell|L \leq \alpha\right) \\
&= P\left(\bigcup_{\ell=1}^{L}\bigcup_{i\in\sigma_\ell}(p_i|\sigma_\ell|L \leq \alpha)\right) \\
&\leq \sum_{\ell=1}^{L} P\left(\bigcup_{i\in\sigma_\ell}(p_i|\sigma_\ell|L \leq \alpha)\right) \\
&\leq \sum_{\ell=1}^{L}\sum_{i\in\sigma_\ell} P\left(p_i|\sigma_\ell|L \leq \alpha\right) \\
&\leq \sum_{\ell=1}^{L}\sum_{i\in\sigma_\ell} \alpha/|\sigma_\ell|L \\
&= \alpha
\end{aligned}
$$

□

Note that when there is a single layer, or when all layers have the same size, the layer-stratified Bonferroni correction is identical to a standard Bonferroni correction.

### 4 An example of the filters used to determine whether a cell population has a true positive significant p-value based on SpecEnr

A significant cell population should: 1) have a mean count of $> 50$ events to prevent inflated ratios, 2) have significantly different actual vs expected proportions for at least one of the sample classes. (if both classes contain actual vs expected proportions that are significantly different, then SpecEnr can be used without filtering), and 3) contains actual and expected proportions that are different at the same rate across both sample classes. For filter 1), we tested thresholds 25, 50, 100, and 200. 50 worked well in practice as we ended up with just as much MDCPs as when we used higher thresholds. However, this threshold may differ based on the data set. For example if one hypothesizes that their MDCP may be a rare cell populations with cell counts of $< 50$ events, then this threshold should be decreased. Note we used a significance threshold of $< .05$ for all T-test p-values.

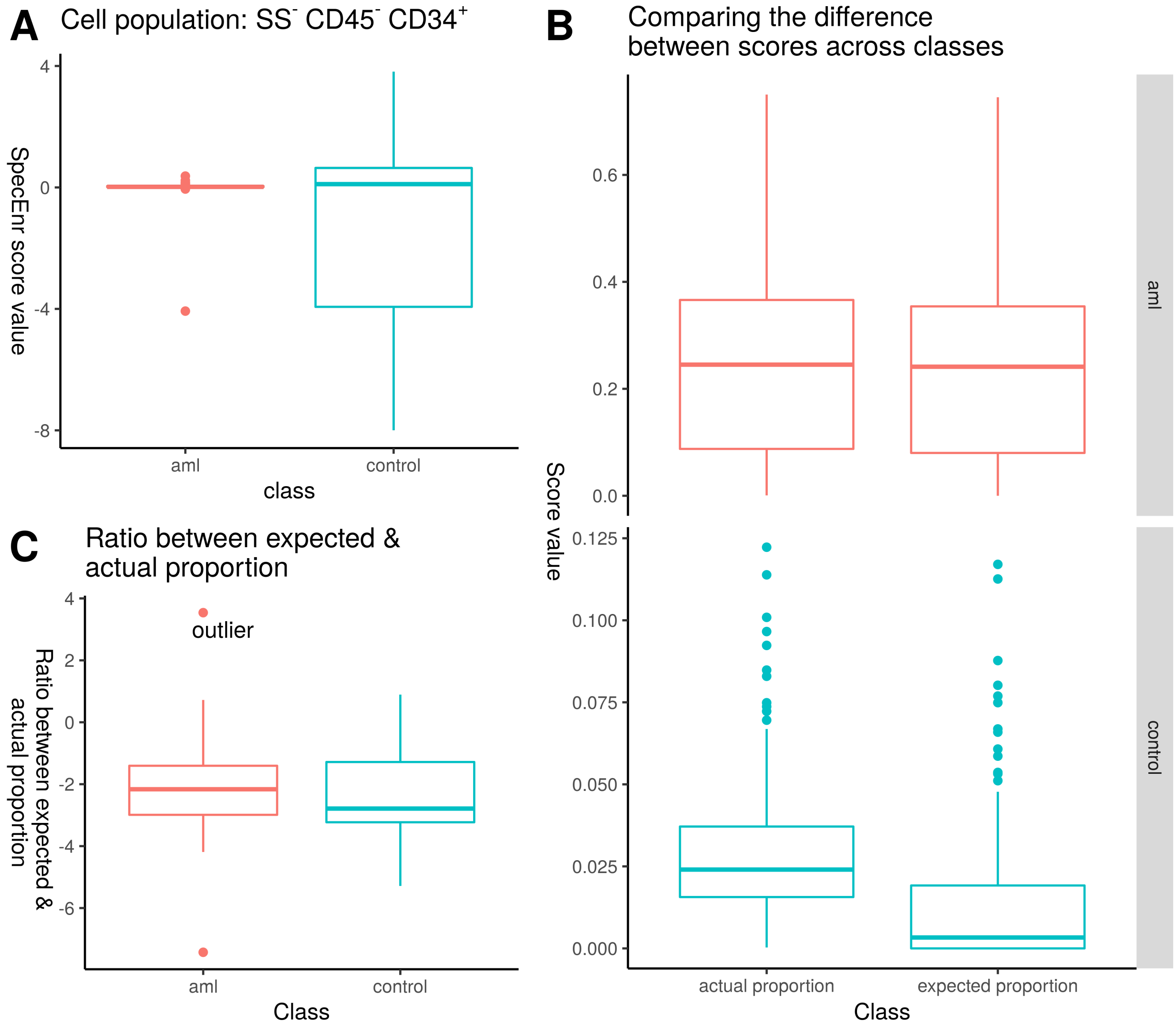


Figure 1: We demonstrate the different filters applied to SpecEnr p-value's using the flowcap data set's $SS^{-}CD45^{-}CD34^{+}$ cell population. A) compares the SpecEnr values between the two classes. B) compares the difference between the actual and expected proportions within sample classes, and C) compares the difference between the actual and expected proportions across sample classes. The comparisons in B) and C) correspond to our three filters.

An example of Filter 2 is shown in Figure 1B where samples of both classes of the flowcap data set have similar actual vs expected proportion values. However, the proportion values from the control class are relatively smaller compare to those of samples from the AML class. To prevent this , we have filter 3), illustrated in Figure 1C. We take the difference between the actual proportion of a random set of 43 control samples and all 43 AML samples. We do the same for expected proportions and we compare these two sets of differences using a T-test.

In Figure 1, cell population $SS^{-}CD45^{-}CD34^{+}$ from the flowcap data set has 1) a mean cell count $> 50$, 2) is significant for the second condition (Figure 1B shows significant and insignificant difference between actual and expected proportions in the control and AML class respectively), and 3) is insignificant for the third condition (Figure 1C shows that the difference between actual and expected proportions across sample classes are not significantly different). Therefore, it is insignificant overall.

### 5 SpecEnr produces robust p-values and q-values

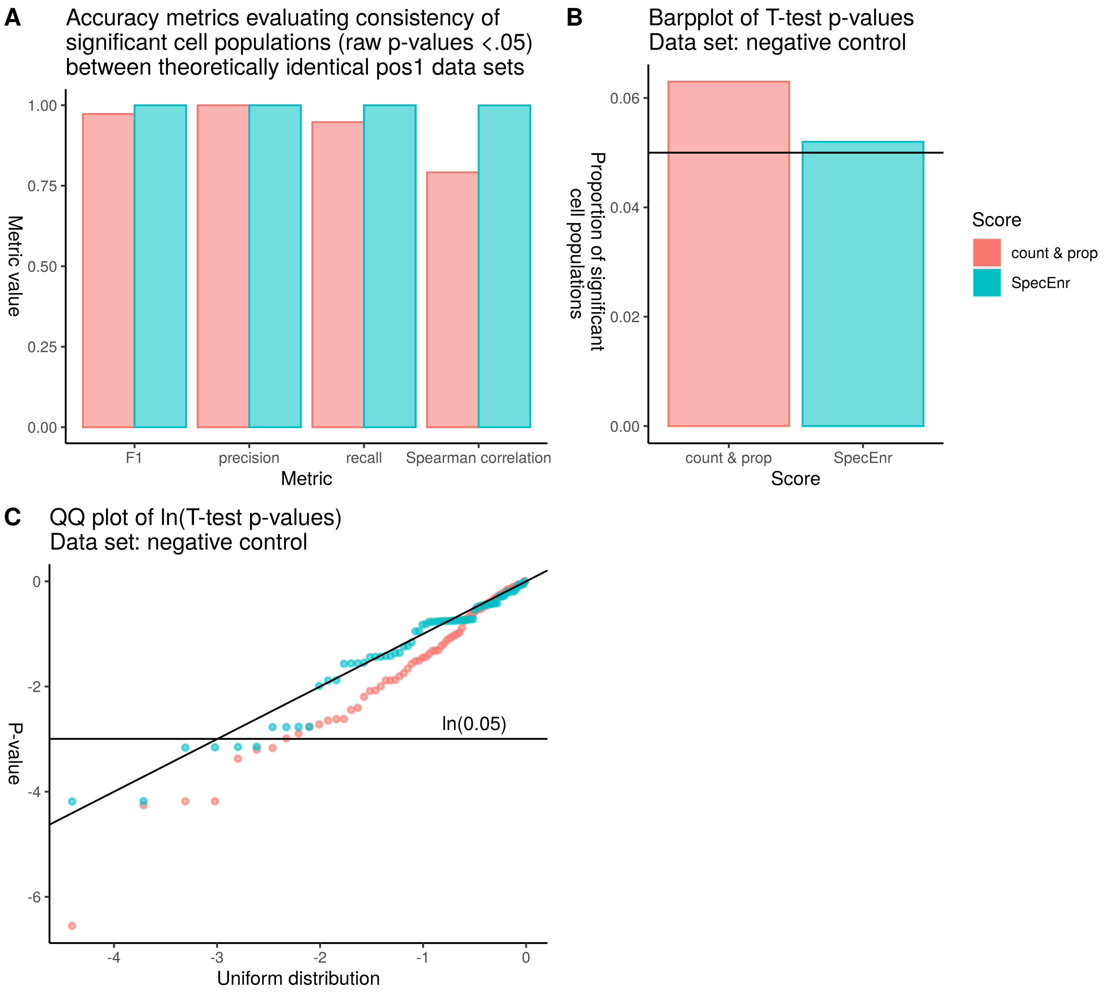


Figure 2: SpecEnr is just as robust as other cell population scores. A) F-measure, recall, precision, and Spearman correlation scores measure high consistency between two theoretically identical positive control data sets (pos1). For the negative control data set (neg1), B) the proportion of cell populations with significant (< .05) p-values is the expected .05, and C) the QQ-plot indicate that unadjusted T-test p-values for all scores align well with a theoretically uniform distribution indicative of our SpecEnr scores' statistical robustness.
.

### 6 Proportion vs SpecEnr plots for data sets pos1-3 where the abundances of the same cell populations are decreased by 50%.

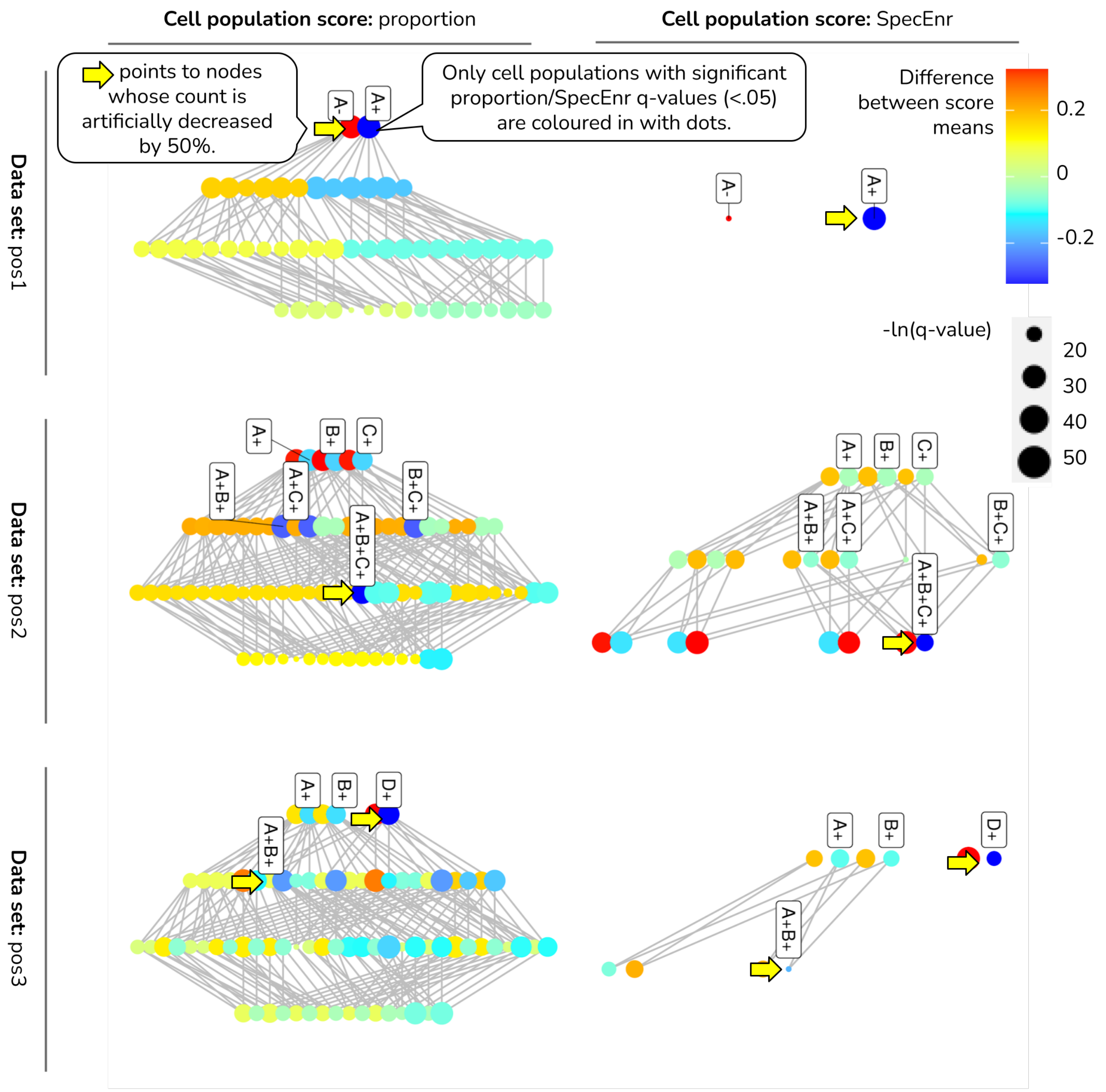


Figure 3: Cell hierarchy plots for the data sets pos1-3, where cell population abundances are decreased instead of increased, indicates that SpecEnr highlights only the MDCPs and their ancestors as opposed to all DCPs as with prop q-values.

### 7 Runtime experiments

Table 1: The runtime in seconds (hours and minutes if the time surpasses a minute) of flowGraph for calculating the edge list of the cell population hierarchy, SpecEnr scores, and T-test (including the filters) are listed in the first table. We also show runtimes for when we use 15 cores vs just 1 core for the pregnancy data set to gauge at potential runtime decreases with hardware improvements. Note that this is not done for the other data sets as setting up the workers for parallel processing would take a few seconds, which would already surpass the runtime for the actual SpecEnr calculation itself.

| data set | flowGraph (1 core) (seconds) | flowGraph (15 cores) (seconds) | no. of measurements | no. of cell populations |
|---:|:---:|:---:|:---:|:---:|
| pos1 | 0.318 | NA | 3 | 27 |
| pos2 | 0.353 | NA | 3 | 27 |
| pos3 | 0.296 | NA | 3 | 27 |
| flowcap | 12.659 | NA | 7 | 2,193 |
| pregnancy | 7,978 | 2,307 | 13 | 109,192 |

Table 2: To see if we can decrease runtime, we also list runtimes for flowGraphSubset, an alternative to the basic flowGraph constructor in the flowGraph package which calculates the edge list, SpecEnr, and T-test q-values (including the filters) for only the cell populations that have a parent population with a significant SpecEnr q-value. This way, we skip calculating everything for cell populations that do not meet this criterion, thereby saving runtime. The assumption that important MDCPs always have at least one parent population with a significant SpecEnr q-value works for almost all cases. However, if the user wants to test multiple class label sets on the same samples (e.g. control vs experiment, age, etc.), we still recommend users to use the basic flowGraph constructor to calculate the SpecEnr score for all cell populations. So, we recommend users to only use flowGraphSubset IF: 1) the users' data set has more than 10,000 cell populations and you want to speed up your calculation time AND 2) you only have one set of classes you want to test on the SAME SET OF SAMPLES (e.g. control vs experiment).

| data set | flowGraphSubset (1 core) (seconds) | flowGraphSubset (15 cores) (seconds) | no. of measurements | no. of cell populations |
|---:|:---:|:---:|:---:|:---:|
| pos1 | 0.135 | NA | 3 | 27 |
| pos2 | 0.155 | NA | 3 | 27 |
| pos3 | 0.154 | NA | 3 | 27 |
| flowcap | 11.371 | NA | 7 | 688 |
| pregnancy | 6,190 | 958 | 13 | 17,991 |
